# A global dataset for the projected impacts of climate change on four major crops

**DOI:** 10.1101/2021.05.27.444762

**Authors:** Toshihiro Hasegawa, Hitomi Wakatsuki, Hui Ju, Shalika Vyas, Gerald C. Nelson, Aidan Farrell, Delphine Deryng, Francisco Meza, David Makowski

**Author notes:** corresponding author(s): Toshihiro Hasegawa.

## Abstract

Reliable estimates of the impacts of climate change on crop production are critical for assessing the sustainability of food systems. Global, regional, and site-specific crop simulation studies have been conducted for nearly four decades, representing valuable sources of information for climate change impact assessments. However, the wealth of data produced by these studies has not been made publicly available. Here, we develop a global dataset by consolidating previously published meta-analyses and data collected through a new literature search covering recent crop simulations. The new global dataset builds on 8314 simulations from 203 studies published between 1984 and 2020. It contains projected yields of four major crops (maize, rice, soybean, and wheat) in 91 countries under major emission scenarios for the 21st century, with and without adaptation measures, along with geographical coordinates, current temperatures, local and global warming levels. This dataset provides a basis for a comprehensive understanding of the impacts of climate change on crop production and will facilitate the rapidly developing data-driven machine learning applications.

## Background & Summary

Climate change affects many processes of food systems directly and indirectly^1^, but the primary effects often appear in crop production. Projections of crop production under future climate change have been studied since the early 1980s. From the 1990s onward, researchers have used future climate data and crop simulation models to project the impacts of climate change on crop yields under various scenarios^2^. Since then, crop simulation models have been used in hundreds of studies to simulate yields for different crops under a range of climate scenarios and growing conditions^3^. The results have been periodically reviewed and assessed by various national and international organisations, in particular by the Intergovernmental Panel on Climate Change (IPCC) working group II, which provides policy-relevant scientific evidence for the impacts of and adaptation to climate change^3^. Review studies covering the last five IPCC assessment cycles confirm that the overall effects are negative but vary significantly among regions^4,5^.

Before 2010, simulation studies were conducted mainly by individual research groups using different climate models, target years, spatial resolution with local management and cultivar conditions. Since 2010, however, significant efforts have been made to coordinate modelling studies through Agricultural Model Intercomparison and Improvement Project (AgMIP)^6^, which intercompares multiple crop models using the standardised inputs. Early AgMIP activities have successfully disentangled sources of uncertainties in crop yield projections and revealed that yield projections are variable among crop models and that model ensemble mean or median often works better than a single model^7–10^, underpinning the importance of datasets based on multiple crop models.

Data sets including crop model simulations produced by AgMIP were subjected to statistical analysis and the results were used to quantify the impacts of climate change on major crops^11,12^. A versatile tool to aggregate simulated results is already available for global gridded studies^13^ to facilitate access to the data. Besides these coordinated efforts, however, many simulation results are scattered and not readily available for meta-analysis. To deliver policy-relevant quantitative information, we need to develop a shared and well-documented database that can be used to assess the impacts of different climate and adaptation scenarios on crop yields.

Here, we compile and develop a global database for potential use for the IPCC Working Group II assessment, drawing on two data sources. The first source of information is the dataset used in the meta-analysis of Aggarwal, et al.^5^, which includes studies considered in the previous five cycles of IPCC assessments^4,14^. The second source is based on a new literature search of studies published during the sixth IPCC assessment cycle (covering the period 2014-2020) reporting crop simulations produced for several contrasting climate change scenarios. Based on these two sources of information, the combined dataset covers all six cycles of the IPCC assessment and can serve as a strong basis for the analysis from the sixth IPCC assessment onward.

The dataset contains the most relevant variables for evaluating climate change impacts on yields of maize, rice, soybean, and rice for the 21st century. They include geographical coordinates, crop species, CO_2_ emission scenarios, CO_2_ concentrations, current temperature levels, local and global warming degrees, the relative changes in yield as a percentage of the baseline period, and the projected effect size of adaptation options.

## Methods

### Data collection

As shown in the PRISMA diagram (Fig. 1), we used two data sources to develop this dataset. The first source is based on the previous meta-analysis by Aggarwal et al.^5^, which includes studies published before 2016 (Aggarwal-DS, hereafter). This metaanalysis builds on the dataset used for the 5^th^ IPCC assessment report^4,14^ and an additional search through three types of databases; Scientific database (Scopus, Web of Science, CAB Direct, JSTOR, Agricola etc), journals and open access repositories, and institutional Websites (FAO database, AgMIP Database, World Bank, etc.) and Google Scholar. See Aggarwal et al.^5^ for details. Briefly, the search terms used by Aggarwal et al.^5^ include “agriculture” or “crop “or “farm” or “crop yield” or “crop yields” or “farm yields” or “crop productivity” or “agricultural productivity” or “maize” or “rice” or “wheat” and “climate change assessment” or “climate impacts” or “impact assessments” or “climate change impact” or “climate impact” or “effect of climate” or “impact of climate change”. The number of selected papers covering the four major crops is 166. We further screened them according to the availability of local temperature rise and geographical information, and traceability, resulting in 99 studies published between 1984 and 2016.

**Fig. 1.**
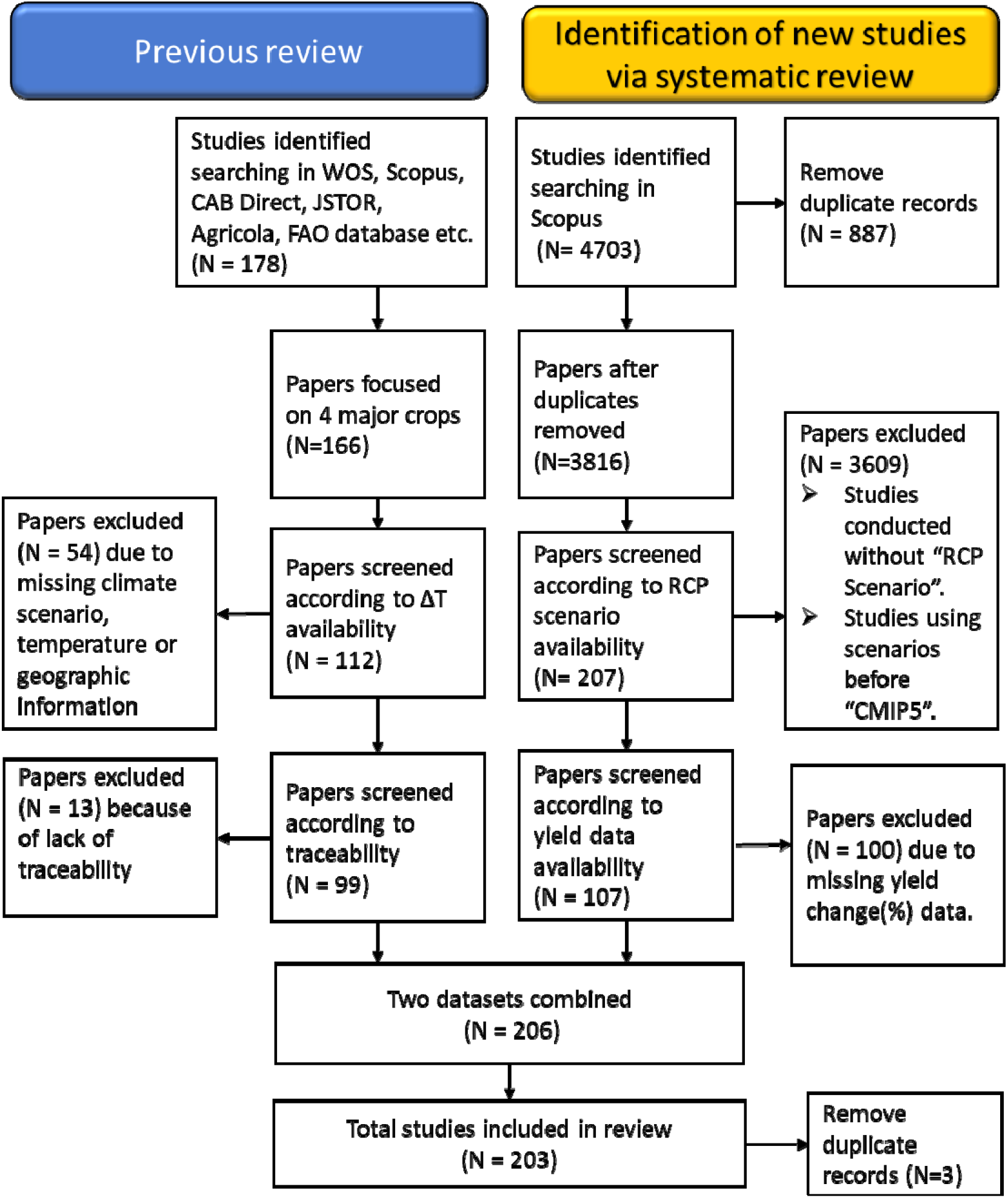
A diagram depicting paper collection and selection from the two data sources. N is the number of studies.

The second source is based on a new recent literature search conducted using Scopus in March 2020 for four major crops (maize, rice, soybean, and wheat) for peer-review papers published from 2014 onward in line with the sixth assessment cycle of IPCC. For maize, the following search equation was used:

PUBYEAR > 2013 AND TITLE-ABS-KEY((maize OR corn) AND ((“greenhouse gas” OR “global warming” OR “climate change” OR “climate variability” OR “climate warming”)) AND NOT (emissions OR mitigation OR REDD OR MRV))

Similar search equations were used for the other crops. Collectively, this search returned a total of 4703 references between 2014 and 2020:1899 for maize, 1790 for wheat, 757 for rice, and 257 for soybean with some duplications because some papers studied multiple crops. Removing the duplicates, the number is down to 3816 studies.

To collect climate-scenario-based simulations, we then selected a subset of studies including the following terms related to climate scenarios in titles, abstracts, or authors’ keywords; “RCP”, “RCP2.6”, “RCP6.0”, “RCP4.5”, “RCP8.5”, “CMIP5”, and “CMIP6”. RCP stands forthe Representative Concentration Pathways^15^, and each RCP corresponds to a greenhouse gas concentration trajectory describing different future greenhouse gas emission levels. The number followed by RCP is the level of radiative forcing (Wm^-2^) reached at the end of the 21^st^ century, which increases with the volume of greenhouse gas emitted to the atmosphere^16^. CMIP5^17^ and CMIP6^18^ are the Coupled Model Intercomparison Project Phase 5 and Phase 6, respectively, where groups of different earth system models (ESMs) provide global-scale climate projections based on different RCPs. Additionally, “process-based model” was used to search in the authors’ keywords to select for studies that use crop simulation models under CMIP5 or CMIP6 climate scenarios. As of March 2020, no results were found for CMIP6 in any search results.

This screening process resulted in a total of 207 references all together for four major crops. These studies were further evaluated for their variables and data availability; studies not reporting yield data were excluded. Projected yields with and without adaptations and yields of the baseline period were extracted, along with geographical coordinates, crop species, greenhouse gas emission scenarios, and adaptation options. We also tried to obtain local and global temperature changes and CO_2_ concentrations as much as possible. The authors of the two grid simulation studies provided the aggregated results for countries or regions^19,20^. The results from different ESMs were averaged.

We consolidated these two sources and removed duplicates, resulting in a total of 203 studies. Both datasets include studies with different spatial scales: site-based, regional, and global. Among these, the results from the global gridded crop models were aggregated to country levels, and we focused on top-producing countries, which account for 95% of the world’s production of each commodity as of 2010 (FAOSTAT, http://www.fao.org/faostat/en/, accessed on September 4, 2020). As a result, the dataset contains 8,314 sets of yield projections during the 21^st^ century from studies published between 1984 and 2020 (Online-only Table 1).

**Table 1.** Summary statistics of climate change impacts on four major crops expresses as per decade impact and per degree impact without adaptation.

|  | Per decade impact (% decade<sup>-1</sup>) |  |  |  | Per degree impact (% °C<sup>-1</sup>) |  |  |  |
| --- | --- | --- | --- | --- | --- | --- | --- | --- |
|  | Maize | Rice | Soybean | Wheat | Maize | Rice | Soybean | Wheat |
| Minimum | -40.0 | -40.8 | -30.0 | -35.4 | -158.7 | -71.7 | -112.7 | -122.7 |
| Maximum | 14.2 | 26.2 | 13.8 | 21.2 | 47.4 | 118.9 | 40.4 | 64.5 |
| Mean | -4.4 | -1.3 | -3.9 | -2.1 | -15.5 | -2.5 | -13.2 | -6.7 |
| 1st quartile | -6.5 | -2.4 | -7.7 | -3.8 | -20.1 | -6.8 | -21.9 | -11.2 |
| Median | -2.2 | -0.7 | -3.3 | -1.2 | -7.3 | -2.2 | -11.7 | -3.6 |
| 3rd quartile | -0.3 | 0.8 | -0.2 | 0.5 | -1.2 | 2.9 | -0.5 | 1.8 |
| Standard deviation | 7.4 | 4.7 | 6.9 | 5.2 | 26.3 | 11.9 | 24.0 | 17.2 |
| Skewness | -1.7 | -2.5 | -0.6 | -1.7 | -2.0 | 0.9 | -1.1 | -2.2 |
| Kurtosis | 7.7 | 21.7 | 4.9 | 10.6 | 7.9 | 23.7 | 5.6 | 13.1 |

### Relative yield impacts

Simulated grain mass per unit land area is used to derive the impact of climate change on yield (YI), which is defined as:

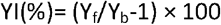

Where Y_f_ is the future yield, and Y_b_ is the baseline yield. However, one study^20^ accounts for technological factors change over time and simulations under both climate change and counterfactual non-climate change scenarios and, for this study, YI is expressed as:

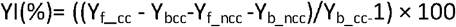

Where Y_f_cc_ and Y_b_cc_ are the future and baseline yield with climate change, Y_f_ncc_ and Y_b_ncc_ is the future yield under counterfactual no climate change scenario.

### Adaptation to climate changes

Various management or cultivar options are tested in the simulations. If the authors of the article consider these options as ways to adapt crops to climate change, we treat them as adaptation options, which are categorised into fertiliser, irrigation, cultivar, soil organic matter management, planting time, tillage, and others. Specifically, in the fertiliser option, if the amount and timing of fertiliser application are changed from the current conventional method, we treat them as adaptation. In the irrigation option, if the simulation program determines the irrigation scheduling based on the crop growth, climatic and soil moisture conditions, we treat this as adaptation because the management is adjusted to the future climatic conditions. If rainfed and irrigated conditions are simulated separately, we do not consider irrigation as an adaptation. We define cultivar option as the use of cultivars of different maturity groups and/or higher heat tolerance compared to conventional cultivars. The planting time option corresponds to a shift of planting time from conventional timing. If multiple planting times are tested, we select the one that gives the best yield. The soil organic matter management option corresponds to application of compost and/or crop residue. The tillage option corresponds to reduced- or no-till cultivation compared to no conventional tillage. When studies consider adaptation options, we compute YI from the ratio of yield with adaptation under climate change to baseline yield without adaptation.

### Warming level

Both local temperature rise (ΔT_I_) and global mean temperature rise (ΔT_g_) from the baseline period have important implications. The former directly affects crop growth and yield, and the latter represents a global target associated with the mitigation activities. We extracted both ΔT_I_ and ΔT_g_ from the literature as much as possible, but ΔT_g_ is not available in many studies. In such cases, we estimated ΔT_g_ using the Warming Attribution Calculator (http://wlcalc.climateanalytics.org/choices). In the dataset, we provide two estimates for ΔT_g_: one from the current baseline period (2001-2010) and the other from the preindustrial era (1850-1900).

### Current temperature level

Current annual mean temperatures were obtained from the W5E5 dataset^21^, which was compiled to support the bias adjustment of climate input data for the impact assessments performed in Phase 3b of the Inter-Sectoral Impact Model Intercomparison Project (ISIMIP3b, https://www.isimip.org/protocol/3/). The W5E5 dataset includes half-degree grid resolution annual mean temperature data, which we extracted for each simulation site or region using the geographic information.

### CO_2_ concentrations

Several studies report two series of yield simulations obtained using two CO_2_ levels to infer the CO_2_ fertilization effects: one obtained with CO_2_ concentrations fixed at the current levels and the other obtained with increased future CO_2_ concentrations provided by the emission scenario considered. In the dataset, we make this explicit in the following two variables:

1. CO_2_: Binary variable equal to “Yes” if future CO_2_ concentrations from the emission scenarios were used and “No” if the current CO_2_ concentration was used for the yield simulations.
2. CO_2_ ppm: if available, CO_2_ concentration was extracted from the original paper. If not, we calculated it from projected changes in CO_2_ concentrations based on the scenarios and periods studied. CO_2_ concentration data were obtained from https://www.ipcc-data.org/observ/ddc_co2.html for CMIP3 and Meinshausen, et al.^16^ (http://www.pik-potsdam.de/~mmalte/rcps/) for CMIP5.

### Baseline correction

Because baseline periods differed between studies, we corrected YI, ΔT_i_, and ΔT_g_ to the 2001-2010 baseline period by a linear interpolation method following Aggarwal et al^22^. Namely, the impacts YI were first divided by the year gap between the future period midpoint year and the baseline period midpoint year of the original study. The impact per year was then multiplied by the year gap from our reference baseline period midpoint year (2005). The same method was applied to express ΔT_I_ relatively to 2001-2010. We also retain original YI and

## Data Records

### Data availability

All the data are stored in the figshare repository, which are freely available at https://doi.org/10.6084/m9.figshare.14691579.v1, where the following files are uploaded:

1. “Projected_impacts_datasheet.xlsx” includes the final dataset after screening.
2. “Meta-data.xlsx” contains the summary of the dataset, such as the definition and unit of the variables used in “Projected_impacts_datatasheet.xlsx”.
3. “Online_only_summary_table.xlsx” contains data distribution, median, and mean impacts of climate change, presented in the online-only tables.
4. “Supplementary_materials.pdf” contains methods for estimating local temperature rise and summary distribution of climate change impacts on four crop yields.
5. “References.docx” provides a list of references that provided data.

### Coverage of the data

A total of 8252 yield simulations are registered in the consolidated dataset. The number of simulations grows exponentially with publication year: 18 in the 1980s, 179 in 1990s, 724 in 2000s and 7331 in 2010s (Online-only Table 1).

About 83% of the simulations use CMIP5 climate scenarios, and 10% use CMIP3. From CMIP5, RCP2.6, RCP4.5 and RCP8.5 are the most used concentration pathways (Online-only Table 2, Fig. 2a). ΔT_g_ from the baseline period (2001-2010) ranges from 0 to 4.8 °C (0.8 to 5.6 °C from the preindustrial period). Almost all simulations with ΔT_g_ > 3°C use RCP8.5, resulting in a greater ΔTg range under CMIP5 (RCPs) than under previous scenarios (SRES and others).

**Fig. 2.**
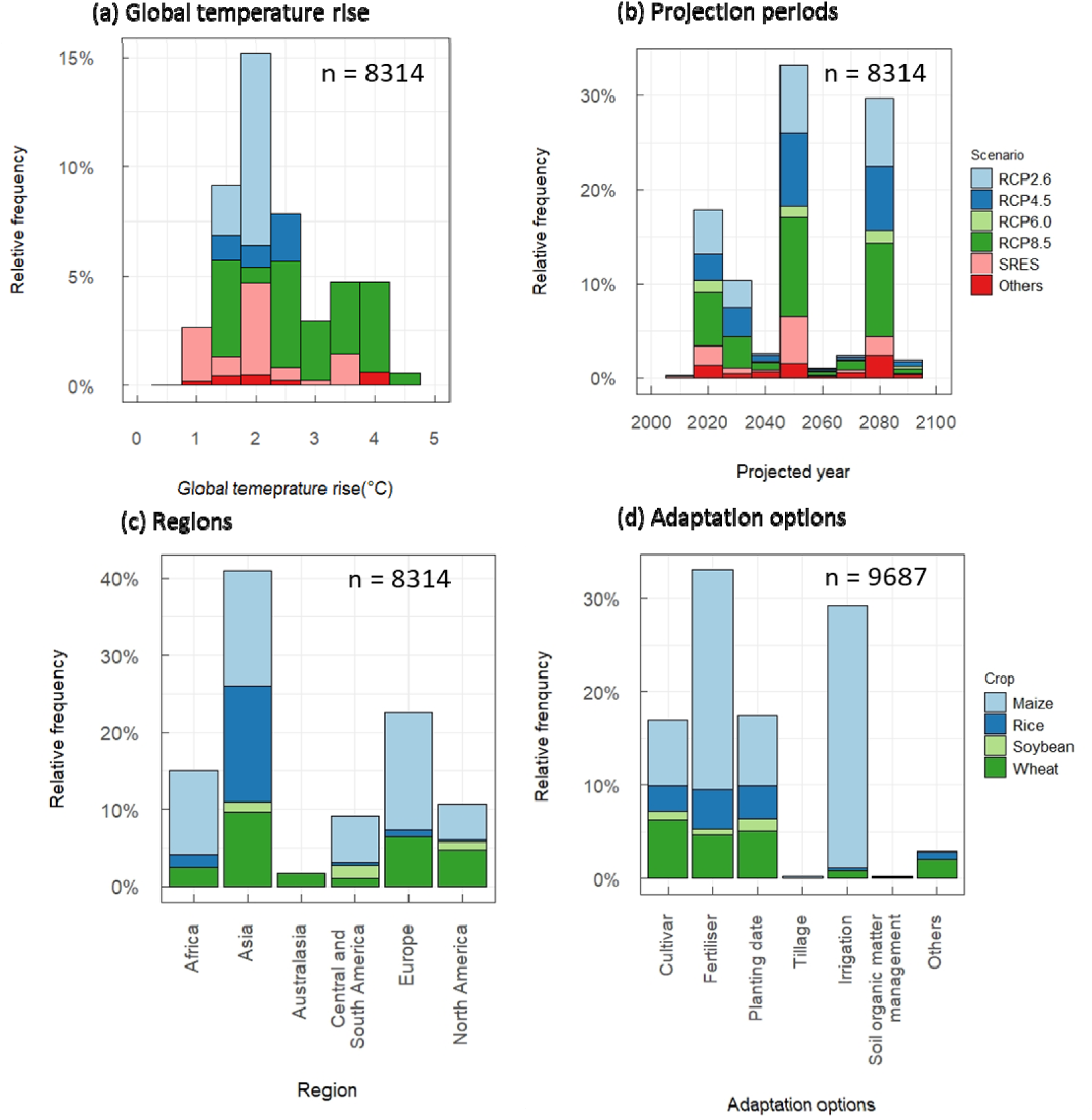
Data availability of crop yield simulations and its breakdown. **(a)** By global temperature rise from the preindustrial era and climate scenarios, **(b)** By projected time periods (midpoint years) and climate scenarios, **(c)** IPCC regions^23^ and crop species, and **(d)** adaptation options and crop species, n is the total number. Note that n=9687 in adaptation options (d) exceeds the total number of simulations (8314) because we collectively add all the options used in the simulations, including those that use multiple options, n is the number of simulation results.

Projected time periods span widely in the 21^st^ century, but the midpoint years peak at 2020 for the near future, 2050 for mid-century, and 2080 for end-century (Fig.2b). Major emission scenarios such as RCP2.6, 4.5 and 8.5 are almost equally distributed across time periods. About 8 % of the simulations assume no CO_2_ fertilisation effects.

Relative frequency of the regions studied generally reflects harvested areas of the four crops in each region (Fig. 2c). About 41% of the simulations were performed in Asia, which accounts for about 47% of the harvested area of the four major crops (mean of 2017-2019, FAOSTAT, http://www.fao.org/faostat/en/, accessed on April 28, 2021). Europe is slightly overstudied for its world share of the harvested areas (12%). Central and South Americas is slightly under-researched (9%) for the regional share of harvested areas (15%), whereas Africa’s share is comparable to the area harvested (10%). Altogether global harvested areas for these four major crops amount 7×10^8^ ha: wheat represents 31% of this area, followed by maize (28%), rice (23%) and soybean (18%). Nevertheless, maize studies are over represented, accounting for about half of the simulations (52%), followed by wheat (26%) and rice (18%); soybean accounts only for 4% of the simulations (Fig.2c). Regionally, maize and wheat are harvested across almost all regions, and simulations follow the actual distribution of these crops. Rice is predominantly studied in Asia, reflecting actual distribution (85 % of the harvested area is in Asia). Soybean remains understudied compared to the other three crops despite its large cultivated area (about 75% of the rice harvested area). Regionally, simulation sites or regions for soybean are mostly in the Americas, which account for 76% of the total soybean harvested area.

About 36 % of the simulations (2992) are performed under current management practices, and the rest (5260) considered different management or cultivars as adaptation options (Fig. 1d). More than half of the simulations are run with multiple options. Among these options, fertiliser accounts for 32 % followed by irrigation (29%), cultivar and planting date (17% each). There are 1853 pairs of yield simulations available for comparing results obtained with and without adaptation. These pairs of yield data can be used to compute the adaptation potentials of the different options considered.

## Technical Validation

### Data quality check

We repeatedly checked the data with multiple authors for the new dataset. For the Aggarwal-DS, we reviewed the sources of references. In case of missing information such as climate scenarios, CO_2_ concentration, or temperature increase, we came back to the original reference. Inconsistencies between the dataset and original papers were corrected when possible. Overall, corrections were made on 163 simulations from 5 studies, which we flag with “*” in the remark column of the dataset. We removed all data of the Aggarwal-DS that were untraceable in the original paper. This quality control excluded 47 simulations from 9 articles listed in the “Excluded” sheet.

We first examined the distribution of the climate change impacts on crop yields, which span from −100 to 136% (Fig. 3). This distribution is skewed to the left, as indicated by the negative skewness. The kurtosis shows that the distribution is salient around the mean compared to the normal distribution. We tested the effects of potential outliers outside the 1.5-fold interquartile range (IQR) on the summary statistics of the climate change impacts on crop yields^24^. Removing values outside the 1.5-fold IQR decreases the number of simulations by 706 (8.5%) and the negative effects of climate change on crop yields by 3.5% for the mean and 0.6% for the median, suggesting that the deletion affected the original distribution. We, therefore, keep all the simulation results in the dataset.

**Fig. 3.**
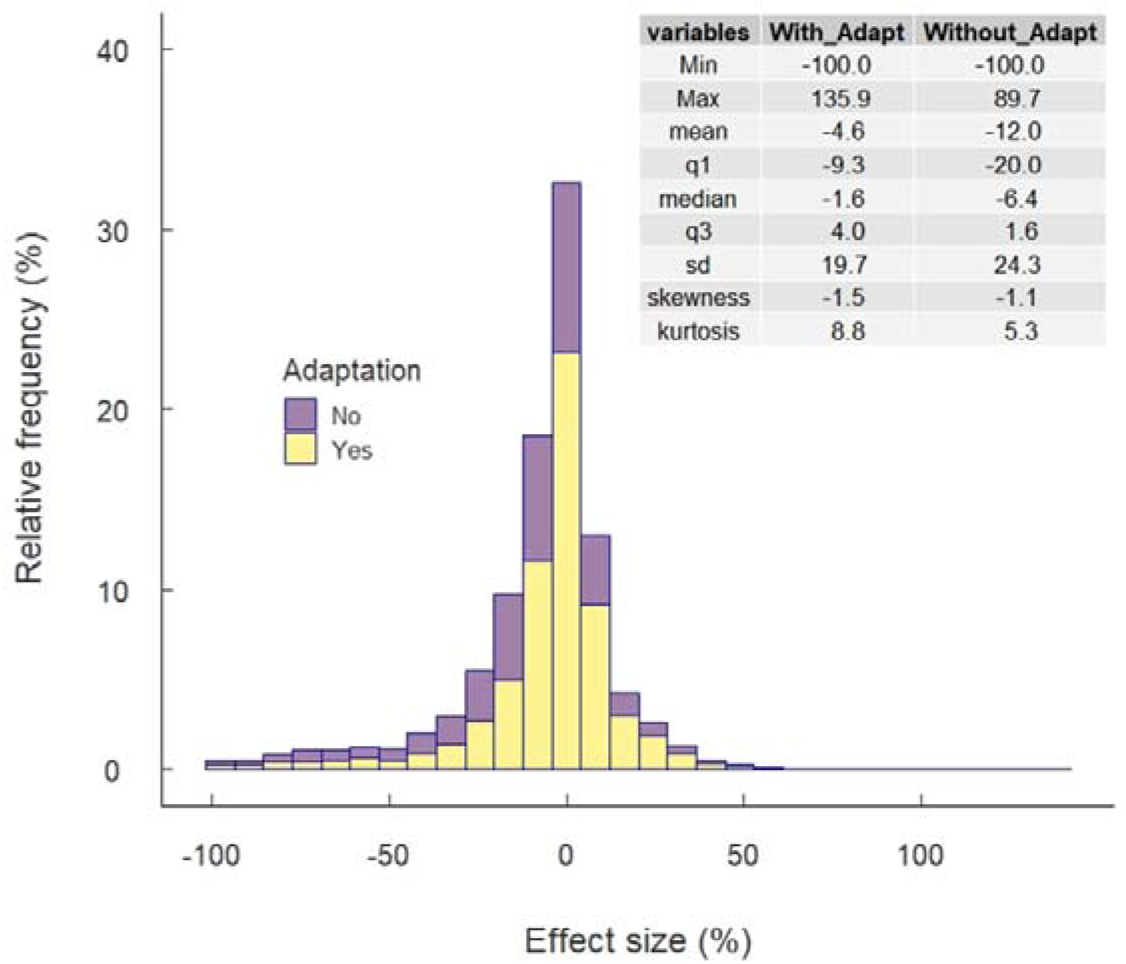
Distribution of climate change impacts on crop yields with and without adaptation

### Methods to estimate local temperature changes

Local temperature change (ΔT_I_) was not available for about half of the simulations. To estimate ΔT_I_ for these studies, we examined the relationship between ΔT_I_ and the following six input variables in 4143 simulations where ΔT_I_ was available: ΔT_g_, CO_2_ concentrations, latitudes, longitudes, time periods, and emission scenarios. Values of ΔT_I_ were estimated using random forest algorithms trained to establish a function relating local temperature rise to the six inputs considered. We tested and compared six models based on different combinations of the input variables. The full model explains about 97.5% of the variation in ΔT_I_, but the reduced model with three variables (ΔT_g_, latitude, and longitude) was close in term of percentage of explained variance (97.0%), and led to a cross-validated RMSE as low as 0.18 °C (Supplementary Table S1 and Fig. S2). We, therefore, used the reduced model to impute ΔT_I_ for the 4110 missing data.

### Comparison with previous studies

The overall effects of climate change on crop yields are negative, with the mean and median of −12% and −6.7% without adaptation and −4.6% and −1.6% with adaptation, respectively (Online-only Table 3 and 4). The median per-decade yield impact without adaptation is −2.3% for maize, −3.3% for soybean, −0.7% for rice, and −1.3% for wheat (Table 1), which are consistent with previous IPCC assessments^14^. The median per-warming-degree impact is −7.6% for maize, −11.7% for soybean, −2.2% for rice, and −3.8% for wheat (Table 2). Per-degree yield impacts for each crop are generally within the range reported in the previous meta-analysis^11^. Among the four crops, soybean has the least number of simulations, resulting in a greater variation in both per-decade and per-degree impacts. Maize consistently shows the largest negative impacts, while rice shows the least.

The climate change impacts by IPCC regional groups^23^ reveals that Europe and North America are expected to be less affected by climate change in the mid-century (MC) and the end-century (EC) than Africa, Central and South America, particularly for maize and soybean. Both positive and negative effects are mixed in all regions (Fig. 4, Supplementary Fig. S2, Fig. S3).

**Fig. 4.**
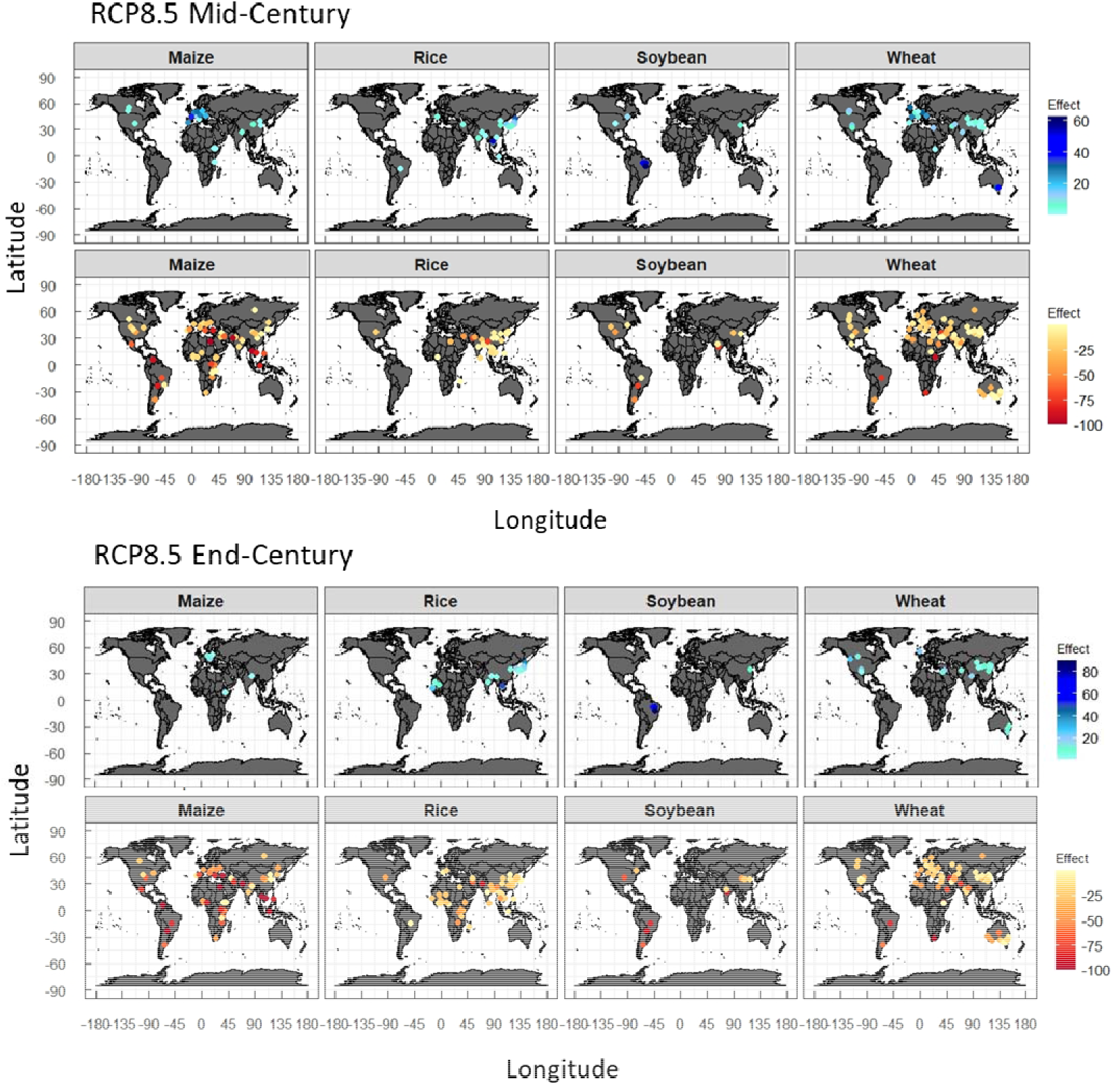
Climate change impacts (% of yield change) on four crops without adaptation under RCP8.5. Upper two panels, Mid-century; Lower two panels, End-Century. Maps with bluish symbols, positive effects (yield gain); Maps with reddish symbols, negative effects (yield loss). Projections under RCP2.6 and RCP4.5 are given in Supplementary Fig. S3.

Regional differences in the impacts in MC and EC are associated with the current temperature level. In MC, positive or neutral effects are projected when current annual average temperatures (T_ave_) are below 10-15 °C, but the effects become negative as T_ave_ increases beyond these levels regardless of RCPs (Fig. 5a). This accounts for the regional differences as a function of latitude reported in previous meta-analyses^4,5^. In EC, the threshold T_ave_ shifts lower, and the negative effects become more severe, particularly under a high emission scenario (RCP8.5) (Fig. 5b). The effect of ΔT_g_ from the baseline period onYI differs depending on the T_ave_ (Fig. 5c); At T_ave_<10°C, YI is generally neutral even where ΔT_g_ >2°C in most crops, but at T_ave_>20°C, YI is negative even with small ΔT_g_, notably in maize. The difference in the YI dependence on ΔT_g_ between regions is also consistent with the previous study^4^.

**Fig. 5.**
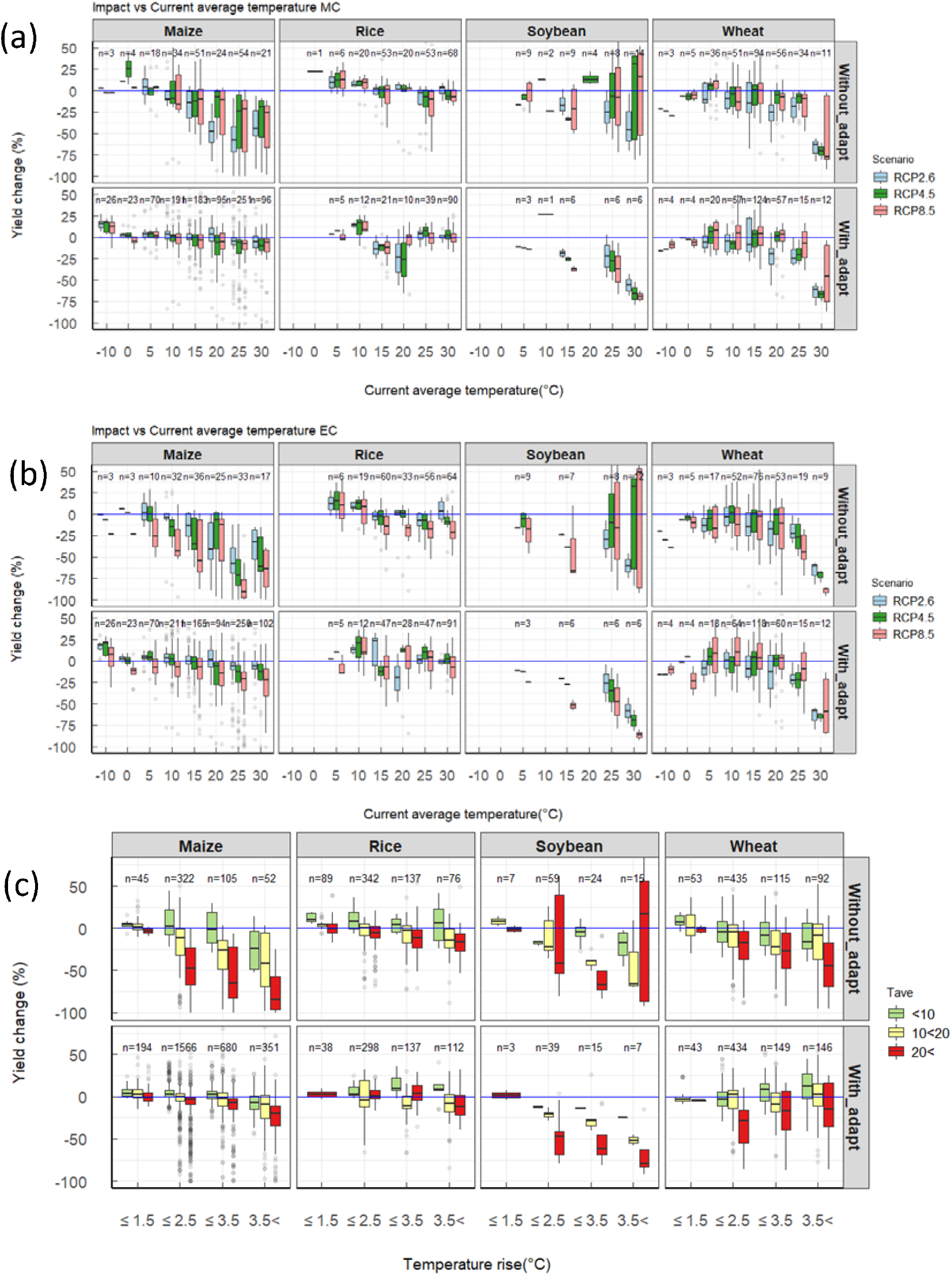
Projected yield changes relative to the baseline period (2001-2005). **(a)** Mid-century (MC) projections without adaptation under RCP8.5 scenario, upper panels showing positive impacts and lower panels negative impacts, (b) End-century (EC) projections under three RCP scenarios by current annual temperature (T_ave_), and **(c)** Yield change as a function of global temperature rise from the pre-industrial period by three T_ave_ levels. The box is the interquartile range (IQR) and the middle line in the box represents the median. The upper- and lower-end of whiskers are median 1.5 × IQR ± median. Open circles are values outside the 1.5 × IQR.

Adaptation potential, defined as the difference between yield impacts with and without adaptation, is an important metric to measure our capacity and limit to adapt to climate change. The options tested with simulation models show an average effect of 8% in MC and 11% in EC (Fig. 6, Supplementary Fig.S4), which is not sufficient to offset the negative impacts, particularly in currently warmer regions. A residual damages will thus likely remain even with adaptation, underpinning several lines of evidence^25,26^.

**Fig. 6.**
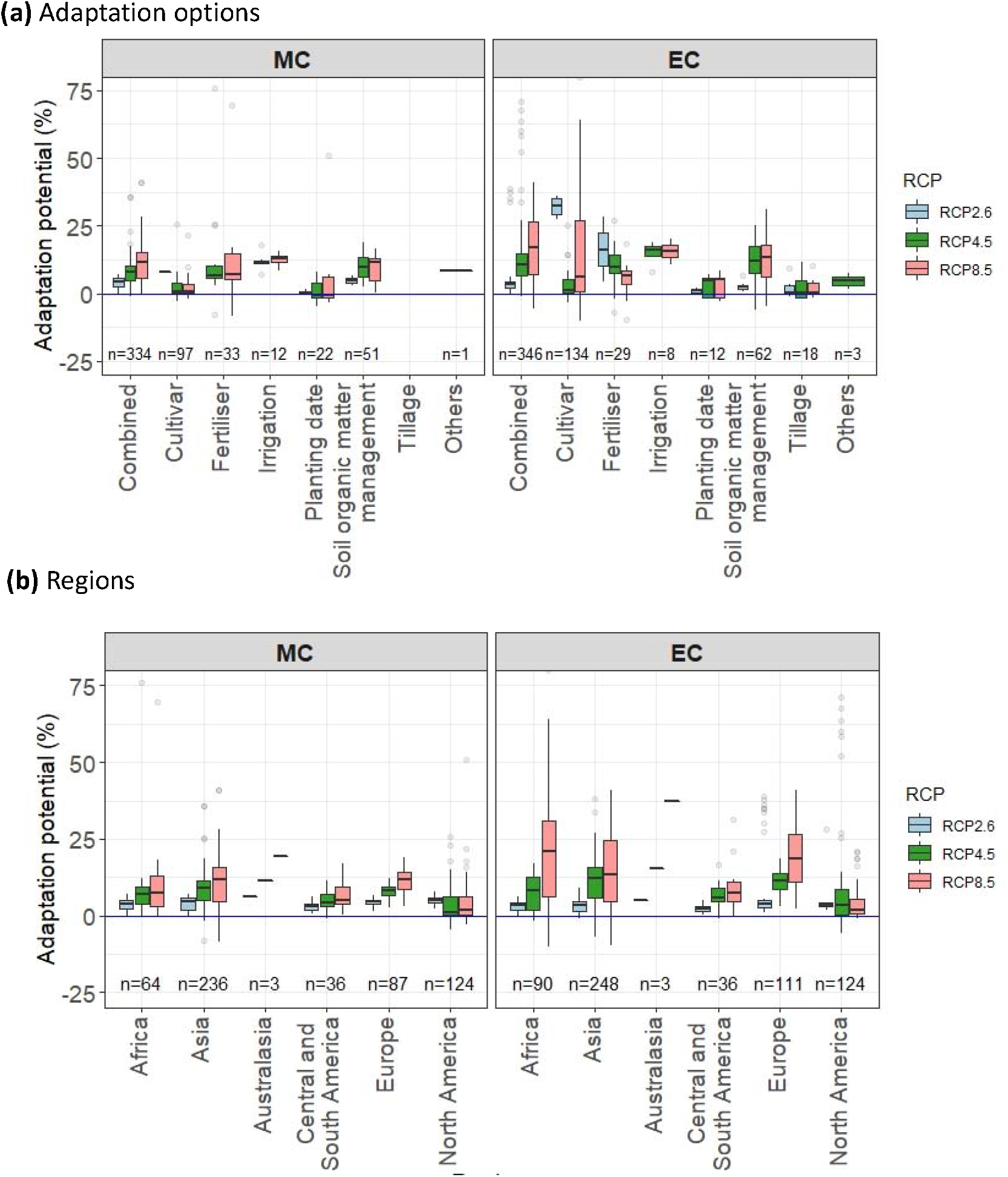
Adaptation potential, defined as the difference between yield impacts with and without adaptation in projected yield impacts, for three RCPs by mid- and end-century (MC, EC). The box is the interquartile range (IQR) and the middle line in the box represents the median. The upper- and lower-end of whiskers are median 1.5 × IQR ± median. Open circles are values outside the 1.5 × IQR. (a) By adaptation options and (b) by IPCC regions.

## Usage Notes

Crop yield simulation studies can provide a narrative of when, where, and what will happen to crop production under different GHG emissions and climate scenarios. They are also expected to provide quantitative information on the potential and limits to adaptation. However, robust estimates covering different temporal and spatial scales need to draw on multiple results obtained from various simulation studies. Nearly four decades have passed since the model projections based on future climate scenarios started. This dataset covers the entire period of simulation studies using climate scenarios, which can help update the quantitative review of climate change impacts on crops. The full list of references is provided in the reference file (https://doi.org/10.6084/m9.figshare.14691579.v1).

Currently, studies are heavily biased towards major cereals such as maize, rice, and wheat, but this can be expanded to include other crops. As of 2020, our literature search failed to find published reports using CMIP6 climate scenarios, but this dataset can be easily updated when new simulations using new climate scenarios or other crop species become available. The next IPCC assessment cycle can fully utilise this dataset by adding the latest simulation results.

One of the caveats to the current dataset is that it only includes crop yield data, notwithstanding crop simulation studies are expected to produce other results than yield. Because of the recent progress in crop modelling, grain quality projections are emerging^27^. We have extensively included the temperature levels to account for the impacts concerning the warming and current temperature, but there is a need to include other key climatic variables such as precipitation and soil moisture. It will be useful to expand our dataset in the future to include this type of data.

## Code Availability

A script file was created using the R statistical programming to estimate ΔT_I_ and is available from the corresponding author upon request.

## Acknowledgements

This study was performed by the Environment Research and Technology Development Fund (JPMEERF20S11820) of the Environmental Restoration and Conservation Agency of Japan. TH and DM would like to thank Joint-Linkage-Call between INRAE and NARO for supporting this collaborative study. We also thank Dr. T. Iizumi and Y. Ishigooka for providing the aggregated simulation results.

## Author contributions

Toshihiro Hasegawa and Hitomi Wakatsuki designed the dataset.

Hitomi Wakatsuki and Hui Ju collected simulation results from the SCOPUS search.

Shalika Vyas designed and collected the Aggarwal dataset.

Gerald C. Nelson conducted literature search and provided global temperature dataset.

David Makowski and Hitomi Wakatsuki developed a statistical imputation for missing data on the local temperature rise.

All authors worked on data analysis and drafting the final version of the manuscript.

## Additional Formatting Information

Correspondence and requests for materials should be addressed to TH.

