## Supplementary information for "A global dataset for the projected impacts of climate change on four major crops"

**Supplementary Table S1.** Comparison of the RF models for estimating local temperature rise as a function of six variables.

| Model | 1 | 2 | 3 | 4 | 5 | 6 |
|---|---|---|---|---|---|---|
| # of variables used | 3 | 3 | 2 | 5 | 4 | 6 |
| Importance of variable | | | | | | |
| Global temperature rise | 1.861 | 0.978 | 1.455 | 0.855 | 1.220 | 1.072 |
| Longitude | 0.362 | X | x | 0.134 | 0.212 | 0.136 |
| Latitude | 0.289 | X | x | 0.195 | 0.293 | 0.231 |
| RCP | x | 0.394 | x | 0.436 | x | 0.309 |
| Future Midpoint year | x | 0.519 | x | 0.503 | x | 0.373 |
| CO2 | x | x | 0.441 | x | 0.566 | 0.327 |
| Fit MSE | 0.033 | 0.142 | 0.118 | 0.058 | 0.035 | 0.028 |
| Fit percent variance explained | 97.07 | 87.29 | 89.43 | 95.04 | 96.92 | 97.52 |
| Median permuted MSE | 0.035 | 0.141 | 0.120 | 0.057 | 0.039 | 0.031 |
| Median permuted percent variance explained | 96.82 | 87.32 | 89.19 | 94.84 | 96.54 | 97.21 |
| Median cross-validation RMSE | 0.181 | 0.375 | 0.348 | 0.231 | 0.191 | 0.174 |

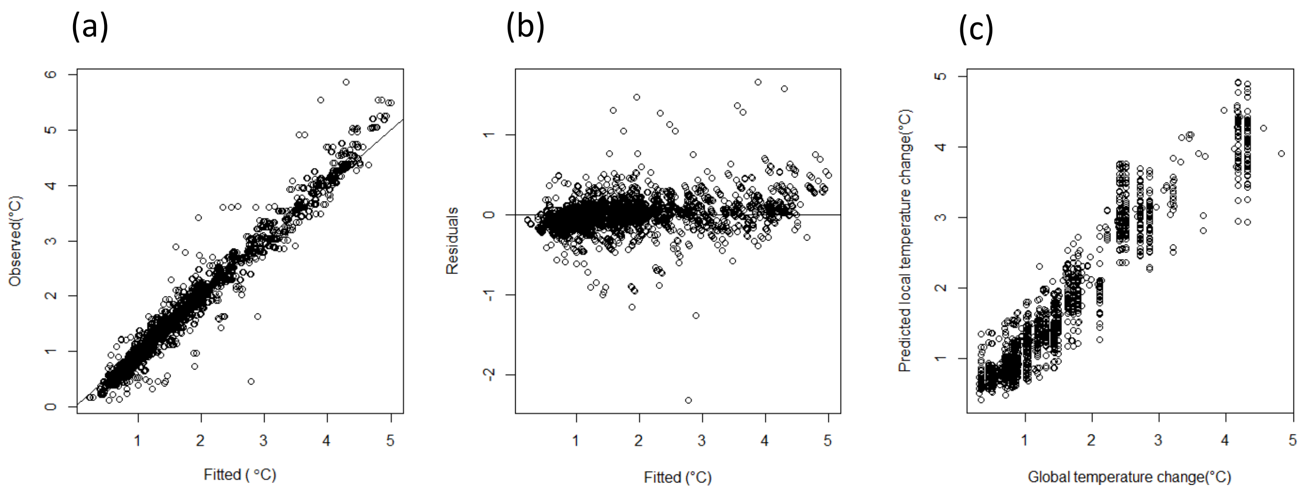

**Supplementary Fig. S1**. Goodness of fit for the Random Forest model for estimating local temperature rise as a function of global temperature rise, latitude and longitude (Model 1 in Table S1). (a) Observed and fitted, (b) Residual plot, (c) Cross-validation prediction

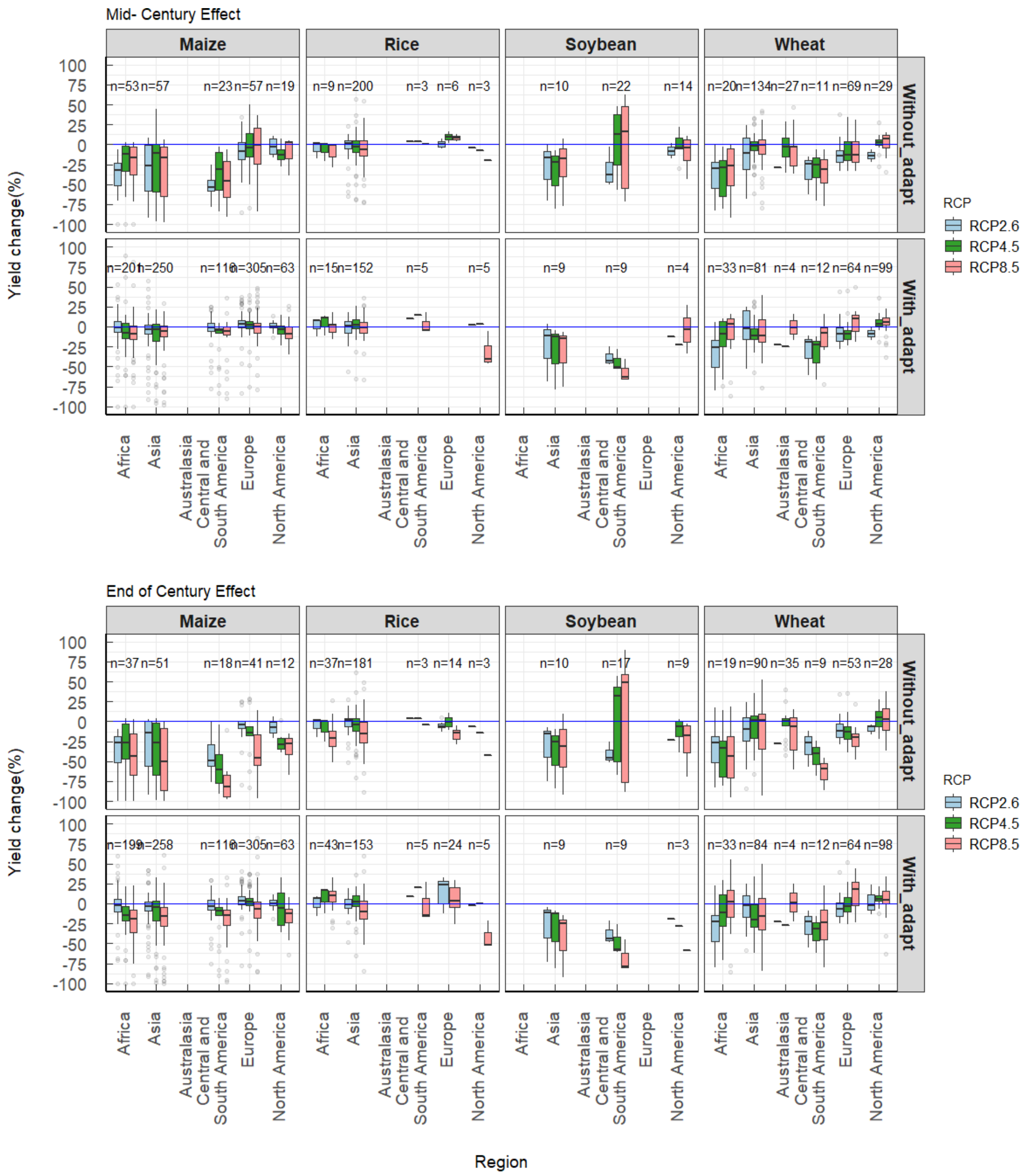

**Supplementary Fig. S2** Climate change impacts on four crops in the mid 21$^{st}$ century with and without adaptation in IPCC regions. Above, mid-century; Below end-century.

RCP2.6 MC No adaptation

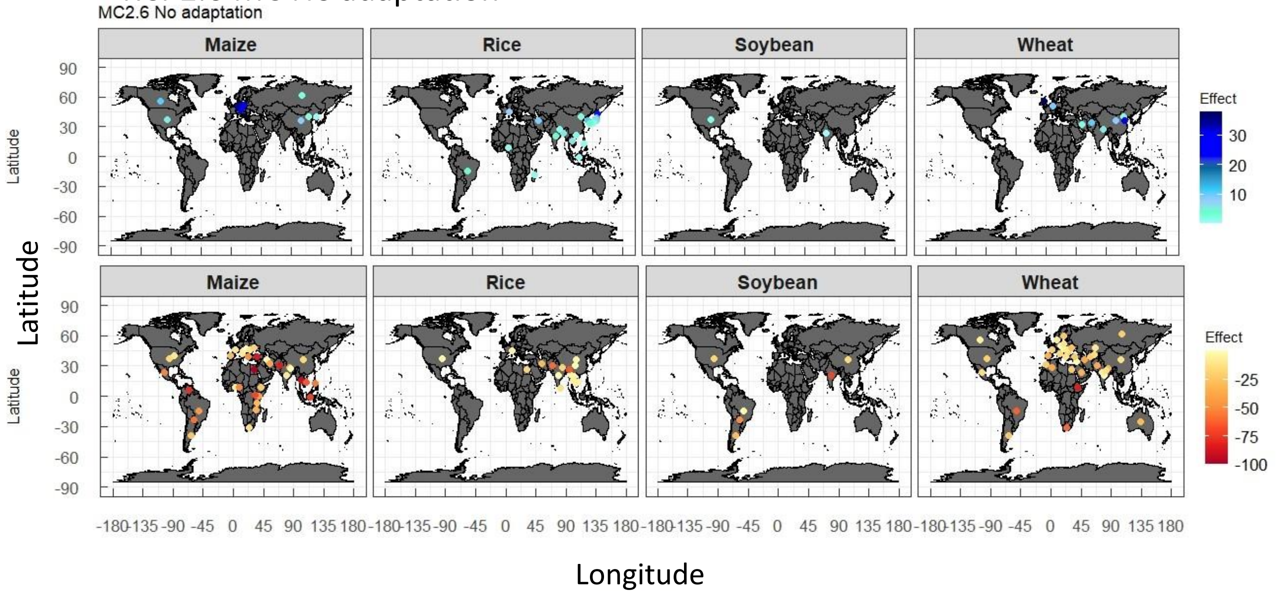

RCP2.6 EC No adaptation

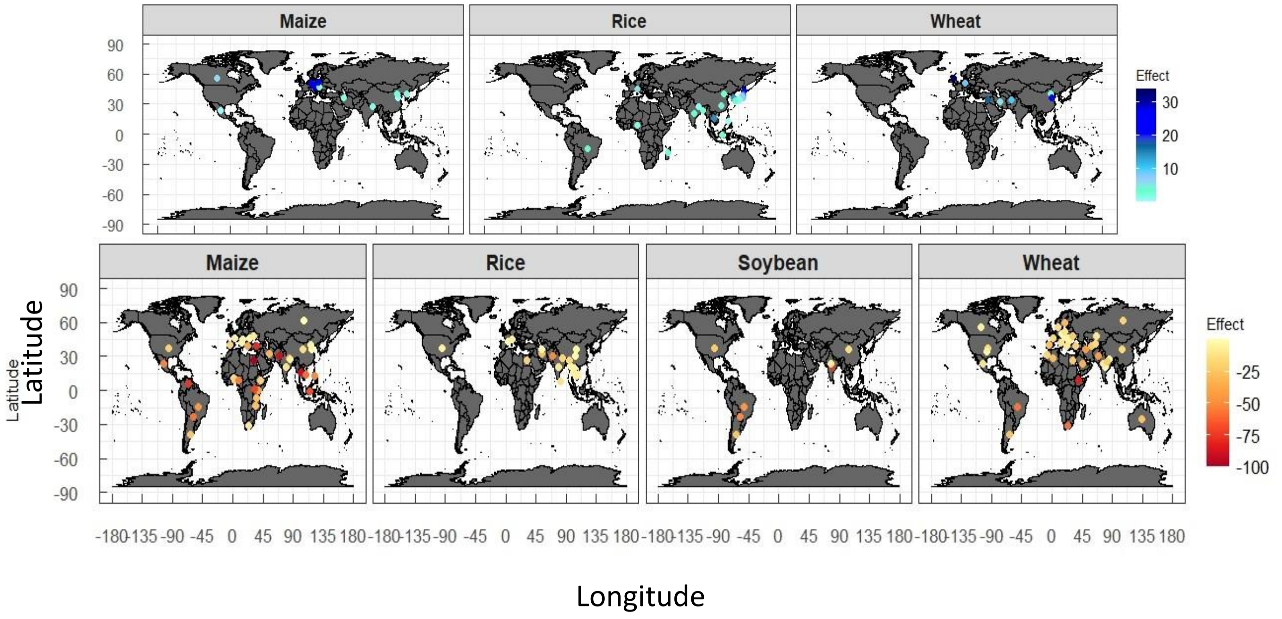

**Supplementary Figure S3(a)** Climate change impacts on four cropswithout adaptation under RCP2.6.

Upper two panels, Mid-century; Lower two panels, End-Century.

Maps with bluish symbols, positive effects; Maps with reddish symbols, negative effects.

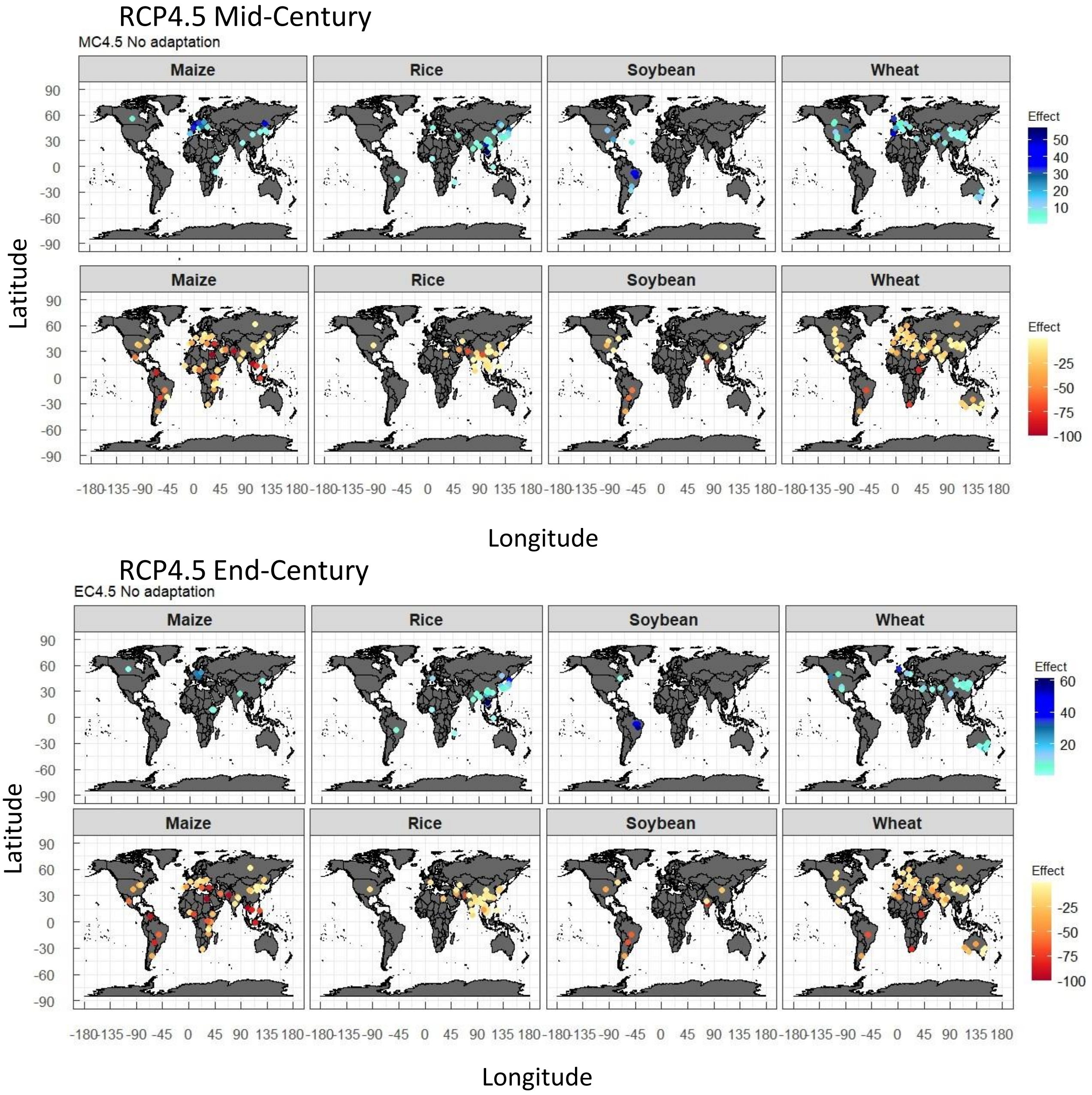

**Supplementary Figure S3(b)** Climate change impacts on four crops without adaptation under RCP4.5.

Upper two panels, Mid-century; Lower two panels, End-Century.

Maps with bluish symbols, positive effects; Maps with reddish symbols, negative effects.

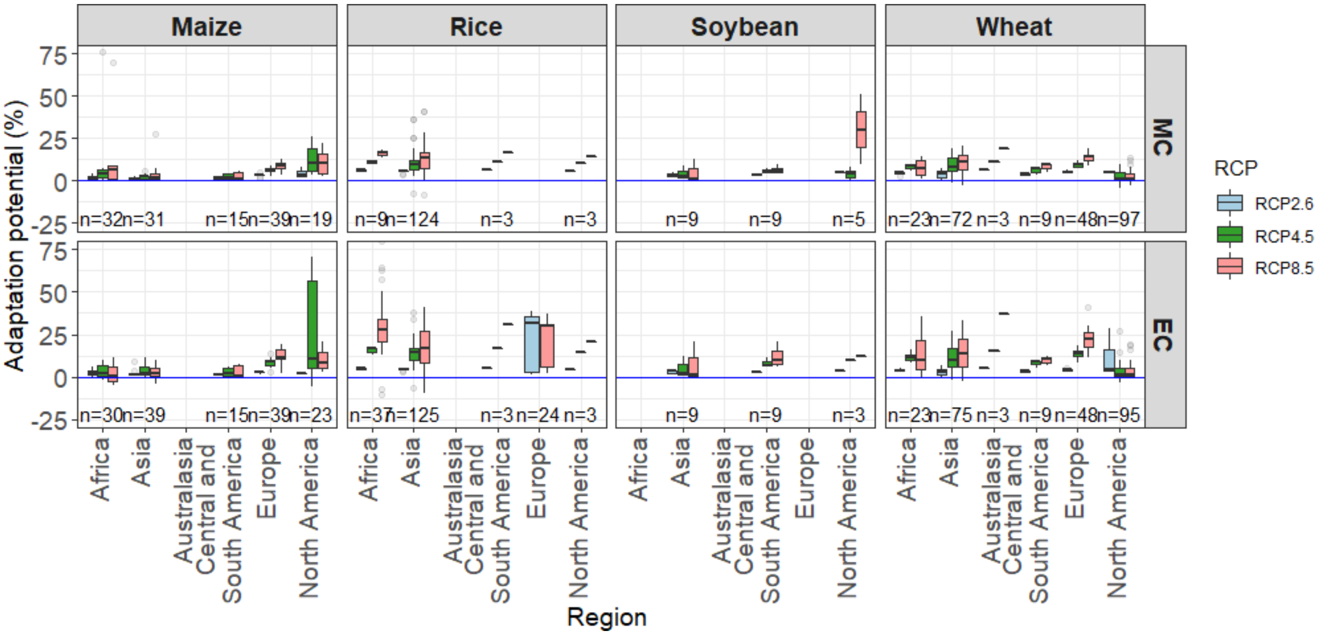

**Supplementary Fig.S4** Adaptation potential in mid- and end century in IPCC regions.
